# Bumblebee queens differ in brain morphology but not learning performance across life stages

**DOI:** 10.1101/2025.10.27.684903

**Authors:** Leeah I Richardson, Shalene Jha, Christopher M Jernigan, Felicity Muth

## Abstract

Animal cognition and brain morphology can vary between individuals and across a lifetime as a function of social and ecological requirements. Bumblebees have distinct ecological pressures acting upon individuals at different times: young queens (gynes) and workers share similar challenges as they both forage as part of the colony, but only queens overwinter and start a colony the following season, switching to a reproductive stage. Here we compared bumblebee (*Bombus impatiens*) visual learning and brain morphology across age-matched workers and gynes and older reproductive queens. We expected foraging-stage gynes to be better at visual learning than reproductive queens and visual regions to be reduced in the reproductive queens, in line with work in other social insects. However, we found that gynes and queens performed similarly, while both performed better than workers. We also found that reproductive queens had larger olfactory regions (antennal lobes) than gynes, while visual regions (medulla and lobula) did not differ, indicating a greater investment in olfactory regions in this later reproductive stage. Brain regions also scaled differently with body size for gynes and workers. Overall, our results provide behavioral and morphological evidence that social and ecological roles in a colony shape cognition and neural investment.

## INTRODUCTION

An animal’s cognition reflects both environmental selection pressures and individual experience, and can vary both between individuals and within individuals as their needs change over time (Shettleworth, 2009; Smid and Vet, 2016). For example, desert locusts (*Schistocerca gregaria*) undergo a distinct change in phenotype from a solitary to a gregarious phase, associated with changes in morphology and physiology but also their cognition: solitary forms are capable of aversive taste learning while later gregarious forms are not, despite both being capable of appetitive learning (Simões et al., 2013; Simões et al., 2016). These differences in cognition align with differences in observed feeding behavior: gregarious forms readily consume chemically-defended plants while solitary forms do not. Changes in behavior across an animal’s lifetime can also be associated with neural changes: food-storing black-capped chickadees (*Poecile atricapillus*) show seasonal changes in spatial memory performance and hippocampal volume, a brain region critical for spatial cognition, in line with foraging demands (Sherry and Hoshooley, 2010). Such changes are expected because neural structures are functionally specialized (Mantini et al., 2013; Tanaka et al., 2012) and metabolically costly to maintain (Niven and Laughlin, 2008). As a result, cognition and its neuromorphological underpinnings are expected to dynamically reflect ecological demands both between and within individuals (Gronenberg and Riveros, 2009) coined as ‘neuroecological theory’ (O’Donnell et al., 2014) which has relevance for a wide range of cognitive tasks.

Foraging is a cognitively demanding behavior that can drive cognitive specialization and associated neuromorphology, which may vary within and across species. Past work has shown that frugivorous primates have larger brains than folivores, which is thought to be related to their more cognitively complex foraging requirements (DeCasien et al., 2017), and seed-caching birds that are dependent on spatial memory for retrieving caches have better spatial cognition and larger hippocampi compared to non-seed caching species (Clayton, 1998). Even within a single seed-caching species (*Poecile gambeli*), birds from populations that seed-cache in harsher environments have better spatial memory and larger hippocampi compared to lower elevation populations (Freas et al., 2012). Honeybees and bumblebees have long been used to study foraging behavior because as generalist nectarivores, they are excellent at learning visual and olfactory associations with floral rewards (Chittka and Thomson, 2001) and because their task specialization, where workers (‘foragers’) focus on food collection, allows for cross-caste comparisons (Heinrich, 1979). While much is known about their learning and its neural underpinnings (Gronenberg and Riveros, 2009; Menzel, 2021; Menzel and Giurfa, 2001), foundational work in neuromorphology makes them an especially ideal system for understanding how bee foraging relates to brain morphology. For example, previous work indicates that the medulla and lobula process visual information peripherally (Kenyon, 1897; Paulk and Gronenberg, 2008; Paulk et al., 2009), while the antennal lobe processes olfactory cues peripherally including floral volatiles (Krofczik, 2008) and socially relevant cuticular hydrocarbons (CHCs) (Sharma et al., 2015). In insects, a major brain region that integrates both visual and olfactory information in the central brain and is known to be involved in learning and memory is the mushroom bodies (MBs), with the MB collar primarily receiving visual inputs (Strausfeld et al., 1998) and the MB lip primarily receiving olfactory inputs (Ehmer and Gronenberg, 2002; Gronenberg, 1999; Mobbs, 1982), where tissue volume can be an indicator of sensory investment.

While past work on workers suggests that the bumblebee brain shows plasticity dependent on age and foraging experience (Jones et al., 2013; Riveros and Gronenberg, 2010), much less is known about cognition and neuromorphological investment in one of the most important castes: queens. Queens offer the opportunity to study how cognition and the brain can change with behavioral requirements, since they transition from a foraging stage to a colony-bound reproductive stage after starting a colony. Gynes are produced at the end of the colony lifecycle, where they forage as part of the colony (Alford, 1969; Goulson, 2010); they then mate and are the sole colony members that overwinter. Gynes then emerge in the spring and forage as solitary individuals during nest-searching and initiation. Foraging gynes face similar challenges to foraging workers in terms of needing to learn which flowers offer the best rewards, although gynes are better at learning color associations than workers (Muth, 2021). After they find a nest site and start a colony, the queens cease foraging and enter a fully reproductive phase once their first batch of foraging workers eclose. In this reproductive phase, queens have developed ovaries, produce offspring for the colony, and remain colony-bound in the dark for the rest of their lives (Goulson, 2010). However, it is unclear whether queen cognition and/ or brain morphology changes between the foraging phase and reproductive phase of their lifecycle.

More broadly, ontogenic shifts from free-living to colony-bound reproduction in insects have been shown to coincide with changes in brain investment: in ant queens (*Messor pergandei* and *Pogonomyrmex rugosus*), peripheral visual regions (medulla) shrank once individuals begin nest-founding (Julian and Gronenberg, 2002), and in another species, *Harpegnathos saltator* colony-bound reproductives had smaller visual regions compared to non-reproductive foragers (Gronenberg and Liebig, 1999; Penick et al., 2021). Similarly, paper wasp (Polistinae) nest-bound foundress queens invested less in visual regions relative to olfactory regions compared with workers (O’Donnell et al., 2014) a pattern also found in facultatively eusocial bees (*Exoneura angophorae*) (Tierney et al., 2025). Despite this effort, it is not known if similar differences in investment exist for bumblebee workers, gynes, and reproductive queens and if these ecological differences also impact the cognitive abilities of these animals.

In this study we addressed inter- and intra-caste variation in cognition and neuromorphology in bumblebees. Specifically, we asked whether queen visual learning and neuromorphology differed between a foraging (‘gyne’) and reproductive (‘queen’) stage in line with ecological requirements, i.e. in line with ‘neuroecological theory’ (Gronenberg and Riveros, 2009; O’Donnell et al., 2014). In addition, we compared age- and experience-matched gynes and workers to ask whether these groups differed in learning performance and in the relative size of sensory (peripheral) and sensory integrating (central) brain regions. While gynes have been found to be better at learning than workers (Muth, 2021), it is unclear how queen learning abilities change across their lifetime. It is also currently unknown how gyne and queen brains scale relative to each other and to those of workers. Assuming queen cognition and neuromorphology are associated with their ecological role, we predicted that foraging-stage gynes would perform better at a visual learning task than older, reproductive-stage queens. In addition, we predicted that gynes would show greater investment in visual regions of the brain relative to reproductive queens, specifically in the medulla, lobula, and mushroom body (MB) collar relative to brain regions that primarily process olfactory information (antennal lobes and mushroom body lip). To ask if queens differ from workers, we compared age- and experience-matched gynes to workers. We expected gynes to learn better based on past findings (Muth, 2021). Finally, we compared investment in the sensory and sensory-integrating regions of the brain across vision (medulla, lobula), olfaction (antennal lobe), and two subregions involved in sensory integration and learning in the central brain, one primarily receiving visual inputs (MB collar) and the other olfactory (MB lip), and predicted that they scaled similarly relative to body size across all bee groups (Table 1).

**Table 1:**
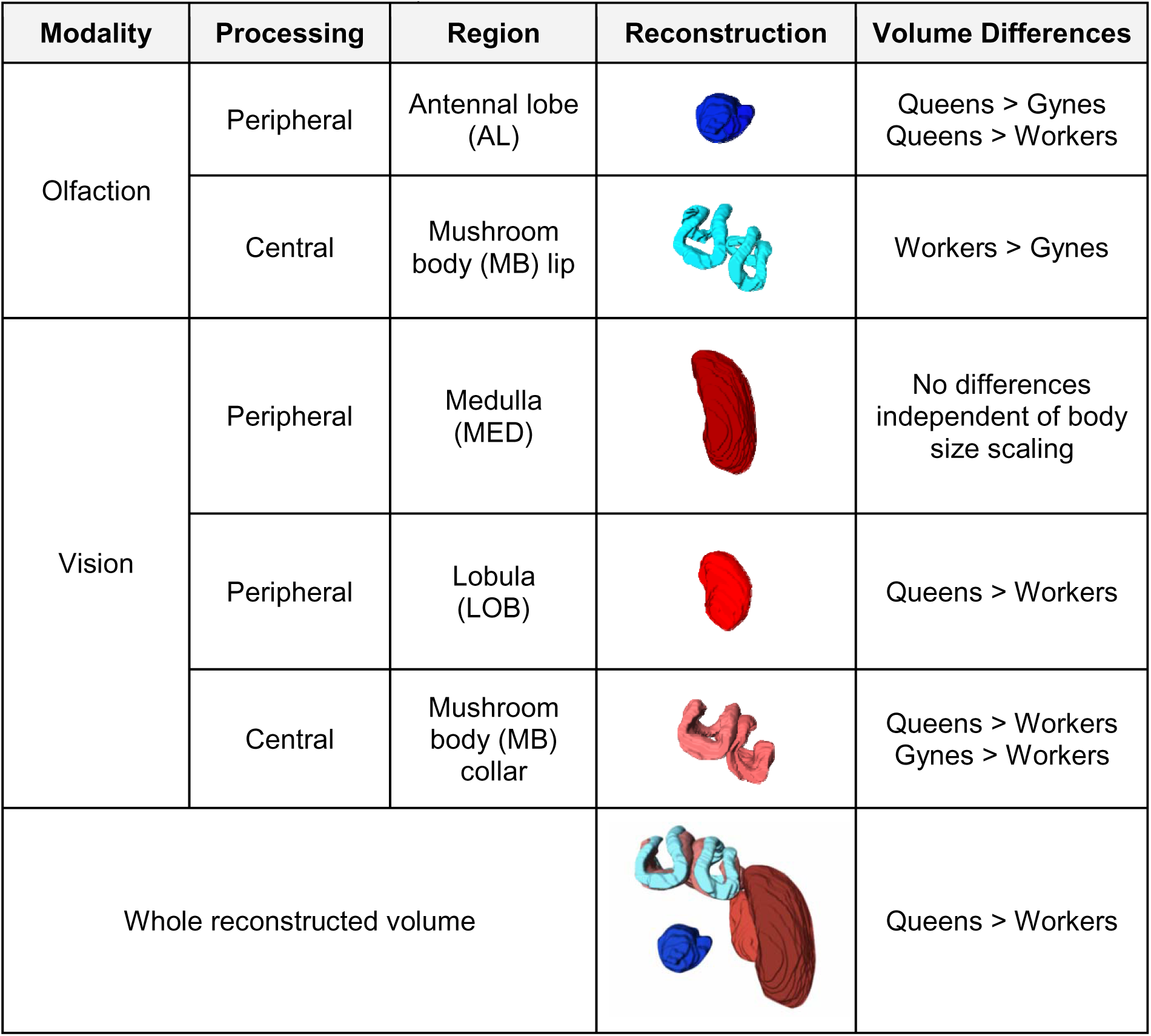
Summary of reconstructed brain regions describing the associated modality (visual vs olfactory) and type of processing (peripheral vs central). Names (including abbreviations) and a 3-D reconstruction of each region are displayed. Summary of the differences in region volume found across groups are also indicated (accounting for body size scaling; significant differences in intercept are shown).

| Modality | Processing | Region | Reconstruction | Volume Differences |
| --- | --- | --- | --- | --- |
| Olfaction | Peripheral | Antennal lobe (AL) |  | Queens > Gynes<br>Queens > Workers |
|  | Central | Mushroom body (MB) lip |  | Workers > Gynes |
| Vision | Peripheral | Medulla (MED) |  | No differences independent of body size scaling |
|  | Peripheral | Lobula (LOB) |  | Queens > Workers |
|  | Central | Mushroom body (MB) collar |  | Queens > Workers<br>Gynes > Workers |
| Whole reconstructed volume |  |  |  | Queens > Workers |

## METHODS

### Colony maintenance and bee selection

We tested 70 bees across 16 commercial bumblebee (*Bombus impatiens*) colonies (Koppert, USA) (Table S1 with sample sizes per colony per assay). Colonies were kept in the boxes that they were received in from Koppert, elevated to allow for airflow and maintained on approximately 3g of honeybee-collected pollen (Koppert, USA) and 30% (w/w) sucrose, replenished 3 times per week. With the exception of when bees were marked for identification, colonies were also kept in dim-light conditions by placing a cardboard lid on top of each colony. We periodically identified all newly eclosed adult bees (workers or gynes) by their whitish unsclerotized appearance and put a small water-based paint mark (Posca, USA) on their thoraces for later identification. Once workers and gynes were 7-10 days old, they were removed from the colony to be tested in a color learning assay (total n = 63; workers = 35; gynes = 28; Table S1). We age-matched workers and gynes to control for effects of age-related changes (i.e. as in Jernigan et al., 2019) as pilot data indicated that age may affect gyne performance (Supplementary Material, Fig. S1). Older queens in a reproductive stage (hereafter ‘queens’) were removed from the colony and underwent the color learning assay once we had finished testing gynes and workers from the same colony, such that all tested bees were taken from a queenright colony (i.e., we did not sample from colonies after the original queen had been removed). This resulted in a total of 7 queens tested; given that we were targeting queens from late-stage colonies that were producing gynes, we did not test queens from the 9 remaining colonies because they died before we were able to test them. As commercially reared queens are unlikely to have free-flying foraging experience including visual foraging experience (Huang et al., 2015), we ensured that all bees did not forage and instead remained in the colony for the duration of their lives. As such, we believe that all bees tested (workers, gynes, and queens) were naïve to visual stimuli and did not have foraging experience. We imaged brains from a subset of bees that underwent the color learning assay (n = 26: workers = 10; gynes = 10; queens = 6; Table S1).

### Color learning

We conducted a free-moving proboscis extension response (FMPER) color discrimination assay to assess differences in visual learning performance between workers, gynes, and queens (first described in Muth et al., 2018, updated in Muth, 2021). We placed bees in testing tubes (2.5cm x 2.5cm x 15cm) for 2 hr 40 min to acclimate and become sufficiently hunger-motivated before training them to discriminate between two shades of human-blue: blue and aqua (same colors as in Muth, 2021, training color counterbalanced across groups; Fig. S3A). Training consisted of three phases: 1) two “no choice” learning trials, 2) four “choice” learning trials, and 3) six unrewarded “test” trials (Fig. 1A). For the no choice trials, we first presented the rewarding color (rewarded conditioned stimulus: CS+) dipped in 50% (w/w) sucrose solution to the bee and allowed her to drink for 3 s, and then presented the unrewarding color (unrewarded conditioned stimulus: CS-) dipped in water and let her drink or antennate for 3 s (bees frequently did not consume the water) (as in Muth, 2021; Muth et al., 2018). For the choice trials, we presented the CS+ and CS- simultaneously such that the bee had to approach and choose one of the stimuli (by extending her proboscis or by antennating it, given that bees can taste via their antennae, Bitterman et al. 1983). After making her choice (denoted as ‘correct’ or ‘incorrect’), we then allowed the bee to sample the other option, so that for each learning trial all bees experienced both the rewarding stimulus and the unrewarding stimulus (Muth, 2021; Muth et al., 2018). We planned the inter-trial interval (ITI) for all six learning trials (2 no choice and 4 choice trials) to be approximately 10 minutes (8-12 min) for all groups; however since the bee ultimately controlled the timing, we measured the actual ITI and found that queens took longer than workers (mean of 12.8 mins vs. 9.2 mins), but that gynes (mean of 10.3 mins) did not differ from workers or queens (X^2^(2) = 7.319, *p =* 0.025; Tukey post-hoc comparisons between groups: queens vs. workers t = 2.566, *p =* 0.033; queens vs gynes: t = −1.717, *p =* 0.207; gynes vs workers: t = 1.472, *p =* 0.311).

**Figure 1.**
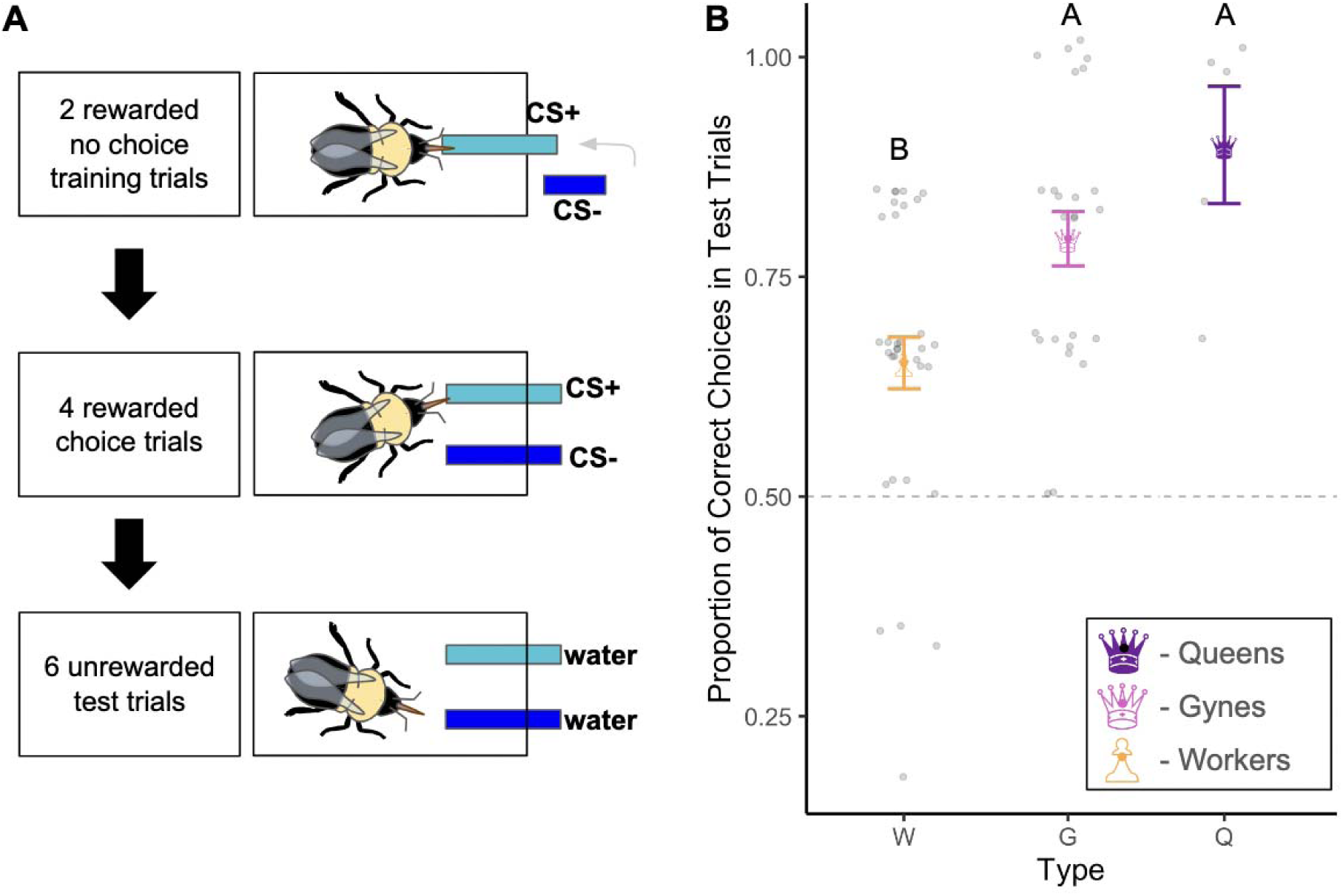
A: Color learning protocol depicting a bee trained to aqua as the rewarding conditioned stimulus (CS+) in three phases. Rewarded stimuli were paired with 50% (w/w) sucrose solution, and unrewarded stimuli (CS-) were paired with water. First, two no choice training trials; second, four rewarded choice trials; finally, 6 unrewarded test trials in which the bee does not make contact with either stimulus. **B:** Performance of each group during the 6 unrewarded test trials, showing significant differences between workers and the other two groups, denoted with letters. Each grey point represents the proportion of correct choices (to the CS+) a single bee made in the test trials; points are jittered for clarity with the group mean +/− standard error depicted in color. The grey dotted line represents performance at chance (50%).

For the unrewarded test trials, the two colors were again presented simultaneously for the bee to choose between, but both strips were dipped in water, and were removed just before the bee contacted the chosen strip; we did this to motivate bees to make multiple choices and to limit learning that strips were unrewarded in the test phase (i.e. extinction learning). These 6 trials were presented such that the bee could make successive choices within a few seconds (i.e. the total test trial lasted an average 5.90 min +/− 6.84 min; mean per group: queens = 8.00 min, gynes = 7.45 min, workers = 4.50 min). If a bee did not move for 12 minutes during the assay, she was presented with a gray strip of paper dipped in sucrose solution and was allowed to drink for 3s to re-motivate her to participate; we excluded bees due to lack of participation if they continued to remain stationary (for 12 mins) after gray strips were presented twice (*sensu* Muth et al., 2021); bees across the three groups did not differ in their motivation to participate in the task (X^2^(2) =-4.425, *p =* 0.109), with 5/7 queens, 25/28 gynes, and 34/35 workers participating. The two blue colors were more chromatically similar to each other than to the gray strips when plotted in bee color space (Fig. S3B), modeled from *B. impatiens* photoreceptor sensitivity (Gawryszewski, 2018; Skorupski and Chittka, 2010) and presentation of gray strips did not affect test trial performance across bee types (Fig. S2C).

For each bee we measured intertegular distance (ITD) (Cane, 1987; Hagen and Dupont, 2013), marginal cell length (Medler, 1962), and hind tibia length (Jernigan et al., 2021) as proxies for body size, by taking photographs under magnification (AmLite, USA) and then measuring each in FIJI (Schindelin et al., 2012). We calculated a PC1 score for each individual from these metrics (Fig. S3) as our measure of body size (as in Jernigan et al., 2021) to account for body size differences when making comparisons between brain region volumes. Body size did not differ between queens and gynes either for those used in the behavioral assays (t(58) = −0.331, *p =* 0.941), or in the subset that underwent brain imaging (t(23) = 0.097, *p =* 0.995). As expected, workers were smaller than gynes and queens in both the full data set (queens vs workers: t(58) = 9.226, *p* < 0.001; gynes vs workers: t(58) = 16.157, *p* < 0.001) and in the subset used for brain imaging (queens vs workers: t(23) = 5.798, *p* < 0.001; gynes vs workers: t(23) = 9.307, *p <* 0.001).

### Dissections and staining

Following participation in the color learning assay, bees that were part of the subset for brain imaging were immediately cold-anaesthetized at 4.4°C (workers: n = 10, although due to damage during processing, one of these brains could only be used for medulla and lobula measurements; gynes: n = 10; queens: n = 6). Once immobilized, bees were decapitated and brains were dissected out of the head capsule in cold PBS (Fig. S4A), before being stored overnight in a 4% paraformaldehyde solution at 4.4°C. We then conducted a staining protocol consisting of three fluorophores in order to visualize neuropils of interest (modified from Jernigan et al., 2019 to include DAPI as a fluorophore in addition to phalloidin TRITC and anti-synapsin Alexa488 and increased incubation times to 5 days to ensure fluorophore penetration in the larger brains; Fig. S4B).

After sitting overnight in the paraformaldehyde fixative, we rinsed the brains twice in PBS, then 6x in PBSTX (1X PBS with 1% Triton X-100 v/v) for 20 min per rinse, and added 1uL of our primary anti-synapsin antibody synorf (mouse anti-fruit fly synapsin, 3C11 DSHB 1:800 dilution) with 10uL of normal donkey serum, and left them to incubate on a shaker for 5 days. We then rinsed the brains 6 times in PBSTX for 20 min each rinse and added 2.5uL of our secondary antibody donkey anti-mouse Alexa488 (Jackson Immuno, USA), along with 10uL of phalloidin conjugated with TRITC (Thermo Fisher, USA), and 1.5uL of the nuclear stain DAPI (Millipore Sigma, USA). Brains were left on the shaker to incubate for an additional 5 days. We then rinsed brains with PBS 6x for 10 min per rinse, followed by a 10 min rinse with 4% paraformaldehyde, and then two more 10 min rinses with PBS. We then conducted an alcohol dehydration, spending 20 min at each concentration in ascending order (30%, 50%, 70%, and 95% ethanol). We kept the brains in 100% ethanol at 4°C for 24 hours and replaced this with another aliquot of 100% ethanol to store in a −20°C freezer prior to imaging.

### Imaging and reconstructions

We cleared the brains by submerging them in methyl salicylate for >15min (up to 1 hour) and conducted whole mount z-stack imaging of one hemisphere with a Zeiss 710 laser scanning confocal microscope with 10X air objective and 12 microns between slices with 16x bidirectional averaging to reduce noise. Imaging was performed at the Center for Biomedical Research Support Microscopy and Imaging Facility at the University of Texas at Austin (RRID:SCR_021756). If one hemisphere did not fit into the frame, two z-stacks were imaged and then tiled together using the 3-D stitching plugin in FIJI (Preibisch et al., 2009; Schindelin et al., 2012). Brain reconstructions were completed by manually tracing neuropils for each region of interest (*sensu* Jernigan et al., 2021) using Amira software (Thermo Fisher, USA).

We reconstructed three peripheral sensory processing brain areas: two visual processing brain regions (medulla and lobula) and the primary olfactory processing region of the brain (antennal lobe). We also subdivided two regions of the mushroom body calyces known to integrate information in the central brain, one integrating visual information (mushroom body collar) and one olfactory information (mushroom body lip) (Fig. 2; Table 1).

**Figure 2.**
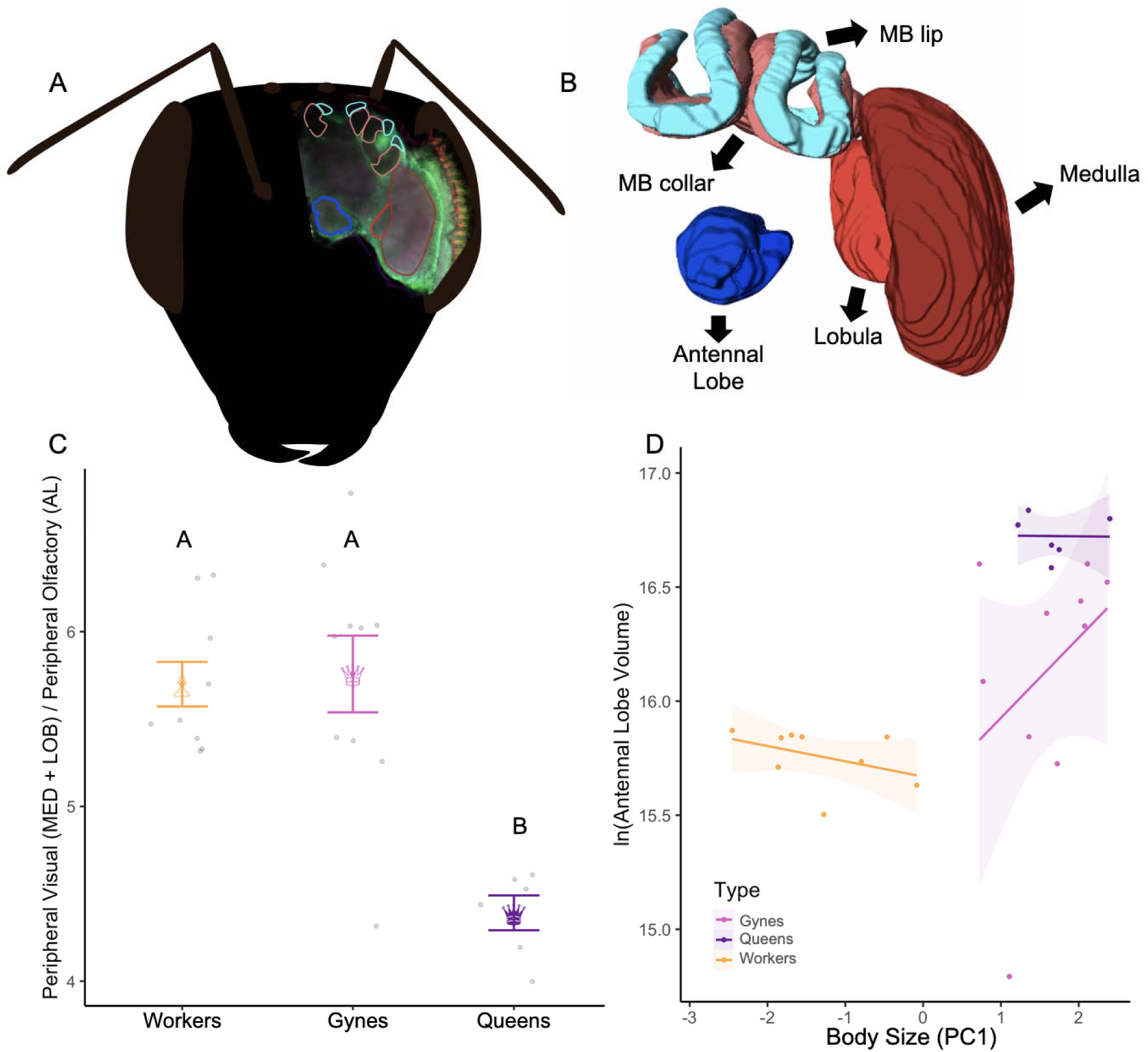
A: Brain regions traced on one slice laid over a bee head to display anatomic relationships **B:** Schematic showing 3D reconstructions - blue tints indicate olfactory regions (dark blue is antennal lobe; light blue is mushroom body lip) and red tints label visual regions (dark red is medulla and red is lobula; light red is mushroom body collar). **C:** Relative investment between peripheral visual regions (medulla and lobula) and the peripheral olfactory region (antennal lobe) showing group means and standard errors. **D:** Natural log of antennal lobe volume (cubic microns) across groups, showing that queens have larger antennal lobes even when accounting for how volume scales with body size; 95% confidence intervals shown around the fitted lines.

### Data analysis

All analyses were run with the R statistical software program (version 4.5.0; R Core Team 2025).

#### Performance across learning trials

To test for the effect of bee type (worker, gyne, or queen) on color discrimination performance during the four learning choice trials, we used a generalized linear mixed model (*glmer*) with a binomial distribution (Bates et al., 2015) with the interaction between bee type (queen, gyne, or worker) and trial number (1-4: continuous) as fixed factors, color the bee was trained to (blue or aqua) as a fixed factor, and individual bee ID as a random factor to account for multiple measures per individual. We determined fixed factor significance with *Anova* in the *car* package (Fox and Weisberg, 2019). For significant terms we used the *emmeans* function in the package *emmeans* (Lenth, 2017) to conduct Tukey posthoc pairwise comparisons within factors.

#### Learning test performance

To test for the effect of bee type (queen, gyne, or worker) on performance in the test phase, we included bee type and the color the bee was trained to as fixed factors and individual bee ID as a random factor. Test trial number was not included in models (i.e. following Muth, 2021; Muth et al., 2018) since test trials were designed to be equivalent (bees’ choices do not change over these 6 trials). We determined fixed factor significance with *Anova* in the *car* package (Fox and Weisberg, 2019). For significant terms we used the *emmeans* function in the package *emmeans* (Lenth, 2017) to conduct Tukey posthoc pairwise comparisons within factors. We also confirmed that each group had learned their trained discrimination (performance above chance at 50%) with a one sample sign-ranked Wilcoxon test for each group, with the alpha value adjusted to 0.017 with a Bonferroni correction due to the three comparisons (Bonferroni, 1936).

#### Brain morphology analyses

To determine if bee types (queen, gyne, or worker) varied in brain region volumes (total constructed volume, medulla, lobula, antennal lobe, mushroom body collar, and mushroom body lip), while accounting for body size scaling (bumblebee worker brain volume scales allometrically (Mares et al., 2005)), we used standard major axis regressions (*sma*) which are commonly used to understand scaling relationships between measures (including work with bees, Cariveau et al., 2016; Yamhure-Ramírez et al., 2025) with bee type (categorical with 3 levels: queen, gyne, or worker) and PC1 of body size (continuous) (similar to Jernigan et al., 2021) as predictors of region volume. These analyses had two parts; first: we tested if region scaled with body size differently for each bee type by comparing slopes. Second: for bee types that had the same slope, we then tested for differences in intercept (Table 1) to assess if their investment in that brain region was predicted by their body size or if they invested relatively more or less (Warton et al., 2012). We used PC1 as our primary measure of body size since it accounts for multiple metrics of body size variation (Jernigan et al., 2021), but our main results hold if we use only inter-tegular distance (ITD) (Table S2), another commonly used body size metric in bee biology (Hagen and Dupont, 2013).

To test for differences in relative investment in visual versus olfactory region brain volumes, we ran two linear models, one for peripheral processing and one for central processing. For peripheral processing we used the ratio between the two relevant visual regions (medulla and lobula) and the antennal lobe (O’Donnell et al., 2014). For central processing we measured the ratio between the mushroom body collar and mushroom body lip (O’Donnell et al., 2014). Relative measures between regions are common in these types of comparisons, and body size was not included as a factor in the models testing differences in relative investment (Jernigan et al., 2021; O’Donnell et al., 2014).

Because the absolute size of brain regions, irrespective of body size, can also be informative of behavioral/cognitive differences (Marino, 2006), we also ran all analyses without accounting for body size to understand absolute differences in brain region volume across our bee types (results shown in Supplementary Material; Table S3).

## RESULTS

### Color learning

All groups of bees learned the color they were trained to, performing above 50% chance level: queens chose the rewarding color 90.0% of the time in the test trials (one-sample sign-ranked Wilcoxon: V = 15, α = 0.017, *p =* 0.002), gynes 79.3% of the time (V = 276, α = 0.017, *p <* 0.001), and workers 65.2% of the time (V = 386, α = 0.017, *p <* 0.001). Queens and gynes made similar numbers of errors in the test phase, but both made fewer errors than workers (*glmer*; full model, type: X^2^(2) = 12.576, *p =* 0.001; color: X^2^(1) = 0.046, *p =* 0.829; post-hoc comparisons: queens vs workers: z = 2.486, *p =* 0.034; gynes vs workers: z = 2.500, *p =* 0.011; queens vs gynes: z = −1.318, *p =* 0.384; Fig. 1B). The three groups of bees did not differ from each other in performance across the learning choice trials (*glmer*; type: X^2^(2) = 0.004, *p =* 0.997), nor did they improve over the trials (*glmer*; trial: X^2^(1) = 0.191, *p =* 0.661; type × trial: X^2^(2) = 0.866, *p =* 0.648; Fig. S5). Bees’ performance also did not vary based on the CS+ color they were trained to (*glmer*; color: X^2^(1) = 2.837, *p =* 0.092; Fig. S2A).

### Neuromorphology

To compare body size-corrected volumes of each brain region across bee types, we first compared the slopes across the three groups to evaluate if the scaling between body size and brain region volume varied by bee type. The total reconstructed volume scaling differed by bee type (*sma*; LRS(2) (likelihood ratio statistic) = 6.860, *p =* 0.032); where gynes’ total reconstructed volume scaled more positively with body size than queens’ (gynes vs queens: t(2) = 4.22, *p =* 0.039) or workers’ (gynes vs workers: t(2) = 5.252, *p =* 0.021), while queens’ did not scale differently with body size compared to workers’ (queens vs workers: t(2) = 0.061, *p =* 0.804). Scaling also differed between bee types for the peripheral brain regions (*sma*; medulla: LRS(2) = 7.704, *p =* 0.021; lobula: LRS(2) = 10.590, *p =* 0.005; and antennal lobe: LRS(2) = 10.53, *p =* 0.005), described further below. Scaling did not differ between bee types for the central brain regions (*sma*; MB collar: LRS(2) = 2.177, *p =* 0.336; MB lip: LRS(2) = 1.900, *p =* 0.386).

For the peripheral visual regions, gyne medullas scaled more positively with body size than worker medullas did (t(2) = 7.484, *p =* 0.006), but there were no other differences in scaling relationships (queens vs workers: t(2) = 0.242, *p =* 0.622; queens vs gynes: t(2) = 2.463, *p =* 0.116). The lobula scaled more positively with body size for gynes than for workers (t(2) = 10.677, *p =* 0.001), but there were no other differences in scaling with body size for the other comparisons (queens vs workers: t(2) = 3.483, *p =* 0.062; queens vs gynes: t(2) = 1.652, *p =* 0.198). Independent of body size scaling (comparing the intercepts), there were no differences in medulla size between queens and gynes (*sma*; intercept: Wald X^2^(1) = 2.96, *p =* 0.085; Table 1), or between queens and workers (*sma*; intercept: Wald X^2^(1) = 0.633, *p =* 0.425; Table 1). Independent of scaling, there was no difference in lobula size between queens and gynes (*sma*; intercept: Wald X^2^(1) = 1.177, *p =* 0.278; Table 1), but queens had relatively larger lobulas than workers (*sma*; intercept: Wald X^2^(1) = 41.910, *p <* 0.001; Table 1).

For the peripheral olfactory region, we found that gyne antennal lobe volume scaled more positively with body size compared to workers (t(2) = 10.528, *p =* 0.001), but there were no other scaling differences between groups (queens vs gynes: t(2) = 1.840, *p =* 0.174; queens vs workers: t(2) = 2.006, *p =* 0.156). Independent of body size scaling, queens had larger antennal lobes than both gynes (*sma*; intercept: Wald X^2^(1) = 8.103, *p =* 0.004; Fig. 2D; Table 1) and workers (*sma*; intercept: Wald X^2^(1) = 83.530, *p <* 0.001; Fig. 2D; Table 1).

For the central processing regions, we found that queens and gynes had larger mushroom body collars, which receives visual input, than workers (*sma*; queens vs workers intercept: Wald X^2^(1) = 19.048, *p <* 0.001; gynes vs workers intercept: Wald X^2^(1) = 5.292, *p =* 0.021; Table 1). Queens and workers did not differ in size of the mushroom body lip, which receives olfactory input (*sma*; queens vs workers intercept: Wald X^2^(1) = 0.968, *p =* 0.325; Table 1), but gynes had a smaller mushroom body lip than workers and marginally smaller than queens when accounting for body size scaling (*sma*; queens vs gynes intercept: Wald X^2^(1) = 3.595, *p =* 0.057; gynes vs workers intercept: Wald X^2^(1) = 4.574, *p =* 0.032; Table 1).

Comparing visual and olfactory region volumes, we found that the relative investment in peripheral visual vs. peripheral olfactory regions varied by bee type (*lm*; F(2) = 15.175, *p <* 0.001; Fig. 2C), with queens investing relatively less in peripheral visual regions (medulla + lobula) relative to the peripheral olfactory region (antennal lobe) compared to gynes and workers (queens vs gynes: t(2) = 7.340, *p <* 0.001; queens vs workers: t(2) = −6.092, *p <* 0.001), and there was no difference between gynes and workers (t(2) = 1.261, *p =* 0.431; Fig 2C). Bee types did not differ in their relative investment of vision and olfaction in central brain areas (*lm*; MB collar / MB lip: F(2) = 2.304, *p =* 0.123).

## DISCUSSION

In this study, we show that queens retain their visual learning capacities between a young foraging and older reproductive stage, while performing better than workers across both stages. In addition to the differences in learning performance across castes, we found differences in brain morphology between both castes and stages: reproductive queens showed a shift in modality-specific peripheral investment evident in their relative investment of vision vs olfaction, distinct from gynes and workers. While we expected visual regions would be greater in gynes relative to reproductive queens, we only found evidence for greater investment in the peripheral olfactory region, suggesting that this region may be especially important for the colony-bound queen. Overall, our results resonate with neuroecological theory, which suggests that ecological demands will be reflected in greater investment in specific brain regions and associated cognitive abilities (Gronenberg and Riveros, 2009; O’Donnell et al., 2014).

First, we did not find that queen visual learning performance was lower than that of gynes despite the behavioral transition to a nest-bound reproductive role. One explanation is that learning capacities are maintained in bumblebee queens to serve a function: colony initiation is turbulent and can be disrupted by abiotic factors such as the length of the winter (Ogilvie and CaraDonna, 2022) or due to other nest-founding queens usurping existing nests (Richards, 1978). The ability to re-initiate nests after early-stage disruptions may vary across species (Tripodi and Strange, 2019) and to our knowledge it is not known if queens are able to re-initiate nests later in colony development. While we expected information processing capacities to be modality-specific (Bitterman, 1965; Frost et al., 2015), and thus visual learning to decline, queens’ learning capacities could also be maintained through domain general processes (Deaner et al., 2006). For example, queens need to maintain dominance hierarchies through aggression in response to worker behavior (van Doorn and Heringa, 1986; Amsalem et al. 2017), monitor brood to make reproductive decisions (Peto et al., 2025), and process social information, as queens are the colony’s social hub, interacting more with daughters than they do with each other (Ruttenberg et al., 2025). It is also possible that older reproductive queens decline in more subtle visual learning capacities than measured here; these could be captured with a more difficult color discrimination or more cognitively complex tasks, such as reversal learning. To control for the lack of foraging experience the older reproductive queens had (since they were commercially reared), we also restricted foraging experience in the gyne and worker groups. However, since foraging experience is related to learning performance (i.e. see Riveros & Gronenberg 2009), it is possible that with foraging experience, differences in learning capacity between queens and gynes would become evident. Nevertheless, previous findings that gynes are better at learning than workers (Muth, 2021) held in our experiment when comparing age-matched naïve individuals, indicating that at least between castes, baseline differences in learning may play a larger role than experience.

Beyond differences in learning, we also found that queens took longer to make choices than gynes and workers, consistent with prior work showing that bumblebee female reproductives are slower decision-makers (Evans and Raine, 2014a). This likely reflects motivational differences, since reproductive queens would not be motivated to forage beyond feeding themselves. It is also possible that some of the differences we see in test trial performance could be explained by speed-accuracy tradeoffs (Chittka et al., 2003). Workers will take risks in terms of sampling previously unrewarding options that could result in finding new floral resources (Evans and Raine, 2014b) so making faster - potentially less accurate - choices may be less costly when foraging as a part of a social colony. Indeed, past work shows that honeybee foragers have two distinct foraging phenotypes which differ in gene expression (Liang et al., 2012) and the concentrations of tyramine and octopamine found in the brain (Cook et al., 2019): the exploratory “scouts” that quickly abandon unrewarding stimuli and tend to prefer novel resources (Carr-Markell and Robinson, 2014; Cook et al., 2020) and the persistent “recruits” that may be more risk-averse (Anselme, 2018). While differences neuromodulator concentrations have not been explored across these two phenotypes in bumblebees, it is possible that similar distinct foraging behavioral repertoires could be relevant to behavioral differences across bumblebee castes. Our findings suggest that reproductive individuals (gynes and queens) may employ different foraging strategies than workers, potentially of importance to their solitary role during colony initiation.

In accordance with our color learning results, we did not detect a reduction in queens’ visual region volume, despite previous work finding this pattern when gynes transition to colony-bound reproductives in ants (*M. pergandei* and *P. rugosus;* Julian and Gronenberg, 2002), and paper wasps (O’Donnell et al., 2014), and reversibly when ants (*H. saltator*) are experimentally manipulated to transition back and forth between colony-bound reproductive and foraging stages (Penick et al., 2021). It is possible that queen brains do not change between the foraging and reproductive stage – however, we found that the olfactory lobe was larger in reproductive queens even when accounting for body size. As such, one possibility for the difference between our results showing no change in visual regions and past studies is that we used naïve individuals. Bumblebee foraging experience is known to increase the size of brain regions relevant to foraging (mushroom body volume) (Riveros & Gronenberg 2010) so it is possible that as gynes begin actively foraging, their brain regions may expand and then later decrease in volume as they transition to reproduction, dependent on experience. Alternatively, there could be general constraints on plasticity of the visual regions within *Bombus impatiens*, as plasticity is known to vary across species (Murren et al., 2015; Touchon et al., 2024) and even between populations of the same species (Gonda et al., 2011; Gonda et al., 2013). For example, in honeybees, mushroom body size increases as workers transition from nurses to foragers, but this change is not reversed if a forager transitions back to a nurse (Fahrbach et al., 2003). It is also possible that the cessation of foraging activity/ flight is energy-saving enough to mitigate the costs of maintaining visual regions since flight increases insect metabolic rate up to 100 fold (Kammer & Heinrich 1978). Past work shows that the effort to learn associations before initiating colonies does not impair queen nest initiation success, even when resources are limited (Watrobska et al., 2024), and our data similarly suggests that maintaining the ability to learn is not costly for queens, though it is possible that longer-term resource limitation could impact maintenance of neural tissue.

In contrast to the results for visual regions, we found that older reproductive queens invested more in peripheral olfaction (antennal lobe volume) than the other two groups, even when body size was accounted for. Olfaction is important for kin recognition in social insects (Van Zweden and d’Ettorre, 2010), and may play a role queen detection of worker-laid eggs (Endler et al., 2004), which queens consume to prevent worker reproduction (Zanette et al., 2012). Olfactory information is likely especially relevant while residing in a dark colony with limited visual information; and visual deprivation is known to cause antennal lobe expansion (Jones et al., 2013). In this study, we assessed differences in learning and differences in brain morphology, but we note that synaptic brain volume is not the only measure of neural investment and organization (Logan et al., 2018) and future work could further explore metrics that may be more sensitive like synaptic density (Kraft et al., 2019), dendritic complexity (Dobrin et al., 2011), or gene expression (Santos et al., 2022).

We also found that queens and gynes showed greater investment in the mushroom body collar (associated with central visual processing, Strausfeld et al., 1998) than workers when accounting for body size scaling. The mushroom body expands as honeybee workers transition into a foraging role (Fahrbach et al., 2003) which is an experience-expectant change that occurs as bees age even in the absence of foraging experience (Fahrbach et al., 1998), although in bumblebee (*B. occidentalis*) workers this expansion appears to be more dependent on experience than age (Riveros and Gronenberg, 2010). The mushroom body is involved in learning and multi-sensory integration (Davis, 1993; Menzel, 2014), and more broadly its volume correlates with learning and sensory sensitivity (Serway et al., 2020). As such, queens’ proportionally larger mushroom body collar region may relate to their greater learning performance, and may reflect an environmental pressure on queens to provision the new colony as the solitary foundress in a visually complex environment.

Overall, we found that queens and gynes performed better than workers in a color discrimination task and invested relatively more in most brain regions (aside from the medulla and lobula) than would be predicted purely by body size. We would also expect gynes and queens to be more sensitive to visual and olfactory information because of their larger size, correlated with eye and antennal size (eyes: Spaethe and Chittka, 2003; Taylor et al., 2019; antennae: Spaethe et al., 2007), which in turn are correlated with visual and olfactory brain region volumes, respectively (visual: Gronenberg and Couvillon, 2010; olfactory: Beltz et al., 2003). Our data supports the idea that, in terms of brain morphology, queens and gynes are not simply larger workers but rather that their brain development is distinct from the worker caste (Cnaani and Hefetz, 2001). Specifically, our finding that queens’ and gynes’ mushroom body collars were disproportionately larger than those of workers’ agrees with broader work on the distributed cognition hypothesis. This hypothesis suggests that larger social insect groups allow for individuals to specialize on tasks, and thus the larger the group size, the smaller individuals’ brains should be (Kverková et al., 2018; O’Donnell et al., 2015). Evidence for this hypothesis is seen in fungus-farming ant species (tribe: Attini), where group size is inversely related to brain size (Riveros et al., 2012), and solitary wasps that have larger mushroom bodies than closely related social species (family: Vespidae) (O’Donnell et al., 2015). Similarly, the larger number of workers within a bumblebee colony may be able to divide tasks and specialize more than queens, which have a solitary phase. In contrast to the distributed cognition hypothesis, the social brain hypothesis suggests that species in larger groups have an increased necessity for larger brains to track interactions and relationships (Dunbar, 1998). While there is substantial evidence for this hypothesis in taxa like primates (Dunbar and Shultz, 2007) there is some support in insects (Lihoreau et al., 2012), including in primitively eusocial bees (Pahlke et al., 2021; Smith et al., 2010) and locusts (Ott and Rogers, 2010). As bumblebee castes are developmentally distinct and cannot be reversed in adulthood (Cnaani et al., 1997), individuals may not need to track social dominance hierarchies, unlike species with more flexible social structures. This caste-based social structure’ and lack of an experience-based dominance hierarchy may explain why distributed cognition is possible in bumblebee colonies.

In summary, our results provide unique support for the bumblebee brain reflecting ecological requirements (i.e. the neuroecological hypothesis). In particular, rather than a reduction in visual brain regions, we found an increase in antennal lobe volume in older reproductive queens, suggesting a shift toward enhanced olfactory processing. This finding is potentially tied to queens’ role within the dark, socially complex nest environment, but could also be related to the lack of visual input all bees experienced. Interestingly, we do not find evidence for increased investment in olfaction necessitating a trade-off with visual regions, since we did not find a reduction in queen visual regions. Additionally, the higher learning performance of gynes and queens relative to workers suggests the possibility of selective pressures on reproductive females to forage efficiently and with minimal risk during the solitary phase. Together, these findings point to a complex relationship between sensory investment and ecological function between bumblebee castes and throughout the lifespan of the queen. Going forward, experimental work could investigate free-foraging bumblebees, sampling at multiple time points throughout their lifecycle to quantify the shifting role of experience (social and foraging) on learning and brain morphology.

## ACKNOWLEDGMENTS

We thank Dr. Wulfila Gronenberg for advice on brain dissection and staining methods, and we thank Paul Oliphint for confocal assistance and troubleshooting.

## REFERENCES

Alford, D. V. (1969). A Study of the Hibernation of Bumblebees (Hymenoptera: Bombidae) in Southern England. J. Anim. Ecol. 38, 149.

Amsalem, E., Padilla, M., Schreiber, P. M., Altman, N. S., Hefetz, A., and Grozinger, C. M. (2017). Do bumble bee, *Bombus impatiens*, queens signal their reproductive status to their workers? Journal of Chemical Ecology. 43(6), 563–572.

Anselme, P. (2018). Uncertainty processing in bees exposed to free choices: Lessons from vertebrates. Psychon. Bull. Rev. 25, 2024–2036.

Bates, D., Maechler, M., Bolker, B. and Walker, S. (2015). Fitting Linear Mixed-Effects Models Using lme4. J. Stat. Softw. 67, 1–48.

Beltz, B. S., Kordas, K., Lee, M. M., Long, J. B., Benton, J. L. and Sandeman, D. C. (2003). Ecological, evolutionary, and functional correlates of sensilla number and glomerular density in the olfactory system of decapod crustaceans. J. Comp. Neurol. 455, 260–269.

Bitterman, M. E. (1965). Phyletic differences in learning. Am. Psychol. 20, 396–410.

Bonferroni, C. (1936). Teoria statistica delle classi e calcolo delle probabilit. Pubblicazioni R Ist. Super. Sci. Econ. E Commer. Firenze 8, 3–36.

Cane, J. H. (1987). Estimation of Bee Size Using Intertegular Span (Apoidea). J. Kans. Entomol. Soc. 60, 145–147.

Cariveau, D. P., Nayak, G. K., Bartomeus, I., Zientek, J., Ascher, J. S., Gibbs, J. and Winfree, R. (2016). The Allometry of Bee Proboscis Length and Its Uses in Ecology. PLOS ONE 11, e0151482.

Carr-Markell, M. K. and Robinson, G. E. (2014). Comparing Reversal-Learning Abilities, Sucrose Responsiveness, and Foraging Experience Between Scout and Non-Scout Honey bee (*Apis mellifera*) Foragers. J. Insect Behav. 27, 736–752.

Chittka, L. and Thomson, J. D. (2001). Cognitive ecology of pollination. Cambridge University Press, UK.

Chittka, L., Dyer, A. G., Bock, F. and Dornhaus, A. (2003). Bees trade off foraging speed for accuracy. Nature 424, 388–388.

Clayton, N. S. (1998). Memory and the hippocampus in food-storing birds: a comparative approach. Neuropharmacology 37, 441–452.

Cnaani, J. and Hefetz, A. (2001). Are queen *Bombus terrestris* giant workers or are workers dwarf queens? Solving the “chicken and egg” problem in a bumblebee species. Naturwissenschaften 88, 85–87.

Cnaani, J., Borst, D. W., Huang, Z.-Y., Robinson, G. E. and Hefetz, A. (1997). Caste Determination in *Bombus terrestris*: Differences in Development and Rates of JH Biosynthesis between Queen and Worker Larvae. J. Insect Physiol. 43, 373–381.

Cook, C. N., Mosqueiro, T., Brent, C. S., Ozturk, C., Gadau, J., Pinter-Wollman, N. and Smith, B. H. (2019). Individual differences in learning and biogenic amine levels influence the behavioural division between foraging honeybee scouts and recruits. J. Anim. Ecol. 88, 236–246.

Cook, C. N., Lemanski, N. J., Mosqueiro, T., Ozturk, C., Gadau, J., Pinter-Wollman, N. and Smith, B. H. (2020). Individual learning phenotypes drive collective behavior. Proc. Natl. Acad. Sci. 117, 17949–17956.

Davis, R. L. (1993). Mushroom bodies and drosophila learning. Neuron 11, 1–14.

Deaner, R. O., van Schaik, C. P. and Johnson, V. (2006). Do Some Taxa Have Better Domain-General Cognition than others? A Meta-Analysis of Nonhuman Primate Studies. Evol. Psychol. 4, 147470490600400114.

DeCasien, A. R., Williams, S. A. and Higham, J. P. (2017). Primate brain size is predicted by diet but not sociality. *Nat*. Ecol. Evol. 1, 0112.

Dobrin, S. E., Herlihy, J. D., Robinson, G. E. and Fahrbach, S. E. (2011). Muscarinic regulation of Kenyon cell dendritic arborizations in adult worker honey bees. Arthropod Struct. Dev. 40, 409–419.

Dunbar, R. I. M. (1998). The social brain hypothesis. Evol. Anthropol. Issues News Rev. 6, 178–190.

Dunbar, R. I. M. and Shultz, S. (2007). Evolution in the Social Brain. Science 317, 1344–1347.

Ehmer, B. and Gronenberg, W. (2002). Segregation of visual input to the mushroom bodies in the honeybee (*Apis mellifera*). J. Comp. Neurol. 451, 362–373.

Endler, A., Liebig, J., Schmitt, T., Parker, J. E., Jones, G. R., Schreier, P. and Hölldobler, B. (2004). Surface hydrocarbons of queen eggs regulate worker reproduction in a social insect. Proc. Natl. Acad. Sci. 101, 2945–2950.

Evans, L. J. and Raine, N. E. (2014a). Changes in Learning and Foraging Behaviour within Developing Bumble Bee (*Bombus terrestris*) Colonies. PLoS ONE 9, e90556.

Evans, L. J. and Raine, N. E. (2014b). Foraging errors play a role in resource exploration by bumble bees (*Bombus terrrestris*). J. Comp. Physiol. A 200, 475–484.

Fahrbach, S. E., Moore, D., Capaldi, E. A., Farris, S. M. and Robinson, G. E. (1998). Experience-Expectant Plasticity in the Mushroom Bodies of the Honeybee. Learn. Mem. 5, 115–123.

Fahrbach, S. E., Farris, S. M., Sullivan, J. P. and Robinson, G. E. (2003). Limits on volume changes in the mushroom bodies of the honey bee brain. J. Neurobiol. 57, 141–151.

Fox, J. and Weisberg, S. (2019). An R companion to applied regression. Sage.

Freas, C. A., LaDage, L. D., Roth, T. C. and Pravosudov, V. V. (2012). Elevation-related differences in memory and the hippocampus in mountain chickadees, *Poecile gambeli*. Anim. Behav. 84, 121–127.

Frost, R., Armstrong, B. C., Siegelman, N. and Christiansen, M. H. (2015). Domain generality versus modality specificity: the paradox of statistical learning. Trends Cogn. Sci. 19, 117–125.

Gawryszewski, F. M. (2018). Color Vision Models: Some simulations, a general n-dimensional model, and the {colourvision} R package. Ecol. Evol. 8(16), 8159–8170.

Godfrey, R. K., Oberski, J. T., Allmark, T., Givens, C., Hernandez-Rivera, J. and Gronenberg, W. (2021). Olfactory System Morphology Suggests Colony Size Drives Trait Evolution in Odorous Ants (Formicidae: Dolichoderinae). Front. Ecol. Evol. 9, 733023.

Gonda, A., Herczeg, G. and Merilä, J. (2011). Population variation in brain size of nine-spined sticklebacks (*Pungitius pungitius*) - local adaptation or environmentally induced variation? BMC Evol. Biol. 11, 1–11.

Gonda, A., Herczeg, G. and Merilä, J. (2013). Evolutionary ecology of intraspecific brain size variation: a review. Ecol. Evol. 3, 2751–2764.

Goulson, D. (2010). Bumblebees: behaviour, ecology, and conservation. Oxford University Press.

Gronenberg, W. (1999). Modality-Specific Segregation of Input to Ant Mushroom Bodies. Brain. Behav. Evol. 54, 85–95.

Gronenberg, W. and Couvillon, M. J. (2010). Brain composition and olfactory learning in honey bees. Neurobiol. Learn. Mem. 93, 435–443.

Gronenberg, W. and Liebig, J. (1999). Smaller Brains and Optic Lobes in Reproductive Workers of the Ant *Harpegnathos*. Naturwissenschaften 86, 343–345.

Gronenberg, W. and Riveros, A. J. (2009). Social Brains and Behavior—Past and Present. In Organization of insect societies: from genome to sociocomplexity, pp. 377–401. Harvard University Press.

Hagen, M. and Dupont, Y. L. (2013). Inter-tegular span and head width as estimators of fresh and dry body mass in bumblebees (*Bombus* spp.). Insectes Sociaux 60, 251–257.

Heinrich, B. (1979). Bumblebee economics. Harvard University Press.

Huang, W.-F., Skyrm, K., Ruiter, R. and Solter, L. (2015). Disease management in commercial bumble bee mass rearing, using production methods, multiplex PCR detection techniques, and regulatory assessment. J. Apic. Res. 54, 516–524.

Jernigan, C. M., Halby, R., Gerkin, R. C., Sinakevitch, I., Locatelli, F. and Smith, B. H. (2019). Experience-dependent tuning of early olfactory processing in the adult honey bee, *Apis mellifera*. J. Exp. Biol. 223, jeb.206748.

Jernigan, C. M., Zaba, N. C. and Sheehan, M. J. (2021). Age and social experience induced plasticity across brain regions of the paper wasp *Polistes fuscatus*. Biol. Lett. 17, 20210073.

Jones, B. M., Leonard, A. S., Papaj, D. R. and Gronenberg, W. (2013). Plasticity of the Worker Bumblebee Brain in Relation to Age and Rearing Environment. Brain. Behav. Evol. 82, 250–261.

Julian, G. E. and Gronenberg, W. (2002). Reduction of Brain Volume Correlates with Behavioral Changes in Queen Ants. Brain. Behav. Evol. 60, 152–164.

Kenyon, F. C. (1897). The Optic Lobes of the Bee’s Brain in the Light of Recent Neurological Methods. Am. Nat. 31, 369–376.

Kraft, N., Spaethe, J., Rössler, W. and Groh, C. (2019). Neuronal Plasticity in the MushroomDBody Calyx of Bumble Bee Workers During Early Adult Development. Dev. Neurobiol. 79, 287–302.

Krofczik, S. (2008). Rapid odor processing in the honeybee antennal lobe network. Front. Comput. Neurosci. 2, 1–13.

Kverková, K., Bělíková, T., Olkowicz, S., Pavelková, Z., O’Riain, M. J., Šumbera, R., Burda, H., Bennett, N. C. and Němec, P. (2018). Sociality does not drive the evolution of large brains in eusocial African mole-rats. Sci. Rep. 8, 1–14.

Lenth, R. V. (2017). emmeans: Estimated Marginal Means, aka Least-Squares Means. 1.10.3.

Liang, Z. S., Nguyen, T., Mattila, H. R., Rodriguez-Zas, S. L., Seeley, T. D. and Robinson, G. E. (2012). Molecular Determinants of Scouting Behavior in Honey Bees. Science 335, 1225–1228.

Lihoreau, M., Latty, T. and Chittka, L. (2012). An Exploration of the Social Brain Hypothesis in Insects. Front. Physiol. 3, 1–7.

Logan, C. J., Avin, S., Boogert, N., Buskell, A., Cross, F. R., Currie, A., Jelbert, S., Lukas, D., Mares, R., Navarrete, A. F., et al. (2018). Beyond brain size: Uncovering the neural correlates of behavioral and cognitive specialization. Comp. Cogn. Behav. Rev. 13, 55–89.

Mantini, D., Corbetta, M., Romani, G. L., Orban, G. A. and Vanduffel, W. (2013). Evolutionarily Novel Functional Networks in the Human Brain? J. Neurosci. 33, 3259–3275.

Mares, S., Ash, L. and Gronenberg, W. (2005). Brain Allometry in Bumblebee and Honey Bee Workers. Brain. Behav. Evol. 66, 50–61.

Marino, L. (2006). Absolute brain size: Did we throw the baby out with the bathwater? Proc. Natl. Acad. Sci. 103, 13563–13564.

Medler, J. T. (1962). Measurements of the Labium and Radial Cell of *Psithyrus* (Hymenoptera: Apidae). Can. Entomol. 94, 444–447.

Menzel, R. (2014). The insect mushroom body, an experience-dependent recoding device. J. Physiol.-Paris 108, 84–95.

Menzel, R. (2021). A short history of studies on intelligence and brain in honeybees. Apidologie 52, 23–34.

Menzel, R. and Giurfa, M. (2001). Cognitive architecture of a mini-brain: the honeybee. Trends Cogn. Sci. 5, 62–71.

Mobbs, P. (1982). The brain of the honeybee *Apis mellifera*. The connections and spatial organization of the mushroom bodies. Philos. Trans. R. Soc. Lond. 298, 309–354.

Murren, C. J., Auld, J. R., Callahan, H., Ghalambor, C. K., Handelsman, C. A., Heskel, M. A., Kingsolver, J. G., Maclean, H. J., Masel, J., Maughan, H., et al. (2015). Constraints on the evolution of phenotypic plasticity: limits and costs of phenotype and plasticity. Heredity 115, 293–301.

Muth, F. (2021). Intra-specific differences in cognition: bumblebee queens learn better than workers. Biol. Lett. 17, 20210280.

Muth, F., Cooper, T. R., Bonilla, R. F. and Leonard, A. S. (2018). A novel protocol for studying bee cognition in the wild. Methods Ecol. Evol. 9, 78–87.

Muth, F., Tripodi, A. D., Bonilla, R., Strange, J. P. and Leonard, A. S. (2021). No sex differences in learning in wild bumblebees. Behav. Ecol. 32, 638–645.

Niven, J. E. and Laughlin, S. B. (2008). Energy limitation as a selective pressure on the evolution of sensory systems. J. Exp. Biol. 211, 1792–1804.

O’Donnell, S., Clifford, M. R., Bulova, S. J., DeLeon, S., Papa, C. and Zahedi, N. (2014). A test of neuroecological predictions using paperwasp caste differences in brain structure (Hymenoptera: Vespidae). Behav. Ecol. Sociobiol. 68, 529–536.

O’Donnell, S., Bulova, S. J., DeLeon, S., Khodak, P., Miller, S. and Sulger, E. (2015). Distributed cognition and social brains: reductions in mushroom body investment accompanied the origins of sociality in wasps (Hymenoptera: Vespidae). Proc. R. Soc. B Biol. Sci. 282, 20150791.

Ogilvie, J. E. and CaraDonna, P. J. (2022). The shifting importance of abiotic and biotic factors across the life cycles of wild pollinators. J. Anim. Ecol. 91, 2412–2423.

Ott, S. R. and Rogers, S. M. (2010). Gregarious desert locusts have substantially larger brains with altered proportions compared with the solitarious phase. Proc. R. Soc. B Biol. Sci. 277, 3087–3096.

Pahlke, S., Seid, M. A., Jaumann, S. and Smith, A. (2021). The Loss of Sociality Is Accompanied by Reduced Neural Investment in Mushroom Body Volume in the Sweat Bee *Augochlora Pura* (Hymenoptera: Halictidae). Ann. Entomol. Soc. Am. 114, 637–642.

Paulk, A. C. and Gronenberg, W. (2008). Higher order visual input to the mushroom bodies in the bee, *Bombus impatiens*. Arthropod Struct. Dev. 37, 443–458.

Paulk, A. C., Dacks, A. M., Phillips-Portillo, J., Fellous, J.-M. and Gronenberg, W. (2009). Visual Processing in the Central Bee Brain. J. Neurosci. 29, 9987–9999.

Penick, C. A., Ghaninia, M., Haight, K. L., Opachaloemphan, C., Yan, H., Reinberg, D. and Liebig, J. (2021). Reversible plasticity in brain size, behaviour and physiology characterizes caste transitions in a socially flexible ant (*Harpegnathos saltator*). Proc. R. Soc. B Biol. Sci. 288, 20210141.

Peto, B. R., Costa, C. P., Moore, M. E. and Woodard, S. H. (2025). Social control of egg-laying in independently nest-founding bumble bee queens. BMC Ecol. Evol. 25, 30.

Preibisch, S., Saalfeld, S. and Tomancak, P. (2009). Globally optimal stitching of tiled 3D microscopic image acquisitions. Bioinformatics 25, 1463–1465.

R Core Team. (2025). R: a language and environment for statistical computing [computer software]. R foundation for Statistical Computing. https://www.R-project.org.

Richards, K. W. (1978). Nest site selection by bumble bees (Hymenoptera: Apidae) in Southern Alberta. Can. Entomol. 110, 301–318.

Riveros, A. J. and Gronenberg, W. (2010). Brain Allometry and Neural Plasticity in the Bumblebee *Bombus occidentalis*. Brain. Behav. Evol. 75, 138–148.

Riveros, A. J., Seid, M. A. and Wcislo, W. T. (2012). Evolution of brain size in class-based societies of fungus-growing ants (Attini). Anim. Behav. 83, 1043–1049.

Ruttenberg, D. M., Wolf, S. W., Webb, A. E., Wyman, E. S., White, M. L., Melo, D., Traniello, I. M. and Kocher, S. D. (2025). Bumble bee workers adopt novel behavioral roles and reshape their social networks in the absence of a queen. 2025.01.07.630106. bioRxiv.

Santos, P. K. F., Galbraith, D. A., Starkey, J. and Amsalem, E. (2022). The effect of the brood and the queen on early gene expression in bumble bee workers’ brains. Sci. Rep. 12, 3018.

Schindelin, J., Arganda-Carreras, I., Frise, E., Kaynig, V., Longair, M., Pietzsch, T., Preibisch, S., Rueden, C., Saalfeld, S., Schmid, B., et al. (2012). Fiji: an open-source platform for biological-image analysis. Nat. Methods 9, 676–682.

Serway, C. N., Dunkelberger, B. S., Del Padre, D., Nolan, N. W. C., Georges, S., Freer, S., Andres, A. J. and De Belle, J. S. (2020). Importin-α2 mediates brain development, learning and memory consolidation in *Drosophila*. J. Neurogenet. 34, 69–82.

Sharma, K. R., Enzmann, B. L., Schmidt, Y., Moore, D., Jones, G. R., Parker, J., Berger, S. L., Reinberg, D., Zwiebel, L. J., Breit, B., et al. (2015). Cuticular Hydrocarbon Pheromones for Social Behavior and Their Coding in the Ant Antenna. Cell Rep. 12, 1261–1271.

Sherry, D. F. and Hoshooley, J. S. (2010). Seasonal hippocampal plasticity in food-storing birds. Philos. Trans. R. Soc. B Biol. Sci. 365, 933–943.

Shettleworth, S. (2009). Cognition, evolution, and behavior. Oxford University Press.

Simões, P. M. V., Niven, J. E. and Ott, S. R. (2013). Phenotypic Transformation Affects Associative Learning in the Desert Locust. Curr. Biol. 23, 2407–2412.

Simões, P. M. V., Ott, S. R. and Niven, J. E. (2016). Environmental Adaptation, Phenotypic Plasticity, and Associative Learning in Insects: The Desert Locust as a Case Study. Integr. Comp. Biol. 56, 914–924.

Skorupski, P. and Chittka, L. (2010). Photoreceptor Spectral Sensitivity in the Bumblebee, *Bombus impatiens* (Hymenoptera: Apidae). PLOS ONE 5, e12049.

Smid, H. M. and Vet, L. E. (2016). The complexity of learning, memory and neural processes in an evolutionary ecological context. Curr. Opin. Insect Sci. 15, 61–69.

Smith, A. R., Seid, M. A., Jiménez, L. C. and Wcislo, W. T. (2010). Socially induced brain development in a facultatively eusocial sweat bee *Megalopta genalis* (Halictidae). Proc. R. Soc. B Biol. Sci. 277, 2157–2163.

Spaethe, J. and Chittka, L. (2003). Interindividual variation of eye optics and single object resolution in bumblebees. J. Exp. Biol. 206, 3447–3453.

Spaethe, J., Brockmann, A., Halbig, C. and Tautz, J. (2007). Size determines antennal sensitivity and behavioral threshold to odors in bumblebee workers. Naturwissenschaften 94, 733–739.

Strausfeld, N. J., Hansen, L., Li, Y., Gomez, R. S. and Ito, K. (1998). Evolution, Discovery, and Interpretations of Arthropod Mushroom Bodies. Learn. Mem. 5, 11–37.

Tanaka, N. K., Endo, K. and Ito, K. (2012). Organization of antennal lobeDassociated neurons in adult *Drosophila melanogaster* brain. J. Comp. Neurol. 520, 4067–4130.

Taylor, G. J., Tichit, P., Schmidt, M. D., Bodey, A. J., Rau, C. and Baird, E. (2019). Bumblebee visual allometry results in locally improved resolution and globally improved sensitivity. eLife 8, 1–32.

Tierney, S. M., Jaumann, S., Hightower, O., and Smith, A. R. (2025). Brain development in a facultatively social allodapine bee aligns with caste but not group living. Frontiers in Ecology and Evolution 13, 1–11.

Touchon, J. C., McMillan, W. O., Ibáñez, R. and Lessios, H. A. (2024). Flexible oviposition behavior enabled the evolution of terrestrial reproduction. Proc. Natl. Acad. Sci. 121, e2312371121.

Tripodi, A. D. and Strange, J. P. (2019). No second chances for pollen-laden queens? Insectes Sociaux 66, 299–304.

van Doorn, A. and Heringa, J. (1986). The ontogeny of a dominance hierarchy in colonies of the Bumblebee *Bombus terrestris* (Hymenoptera, Apidae). Insectes Sociaux 33, 3–25.

Van Zweden, J. S. and d’Ettorre, P. (2010). Nestmate recognition in social insects and the role of hydrocarbons. 222–243. Cambridge University Press.

Warton, D. I., Duursma, R. A., Falster, D. S. and Taskinen, S. (2012). smatr 3 - an R package for estimation and inference about allometric lines. Methods Ecol. Evol. 3, 257–259.

Watrobska, C. M., Šima, P., Ramos Rodrigues, A. and Leadbeater, E. (2024). Potential costs of learning have no detectable impact on reproductive success for bumble bees. Anim. Behav. 214, 173–185.

Yamhure-Ramírez, D., Wainwright, P. C. and Ramírez, S. R. (2025). Sexual dimorphism and morphological integration in the orchid bee brain. Sci. Rep. 15, 8915.

Zanette, L. R. S., Miller, S. D. L., Faria, C. M. A., Almond, E. J., Huggins, T. J., Jordan, W. C. and Bourke, A. F. G. (2012). Reproductive conflict in bumblebees and the evolution of worker policing: Reproductive conflict and policing in bumblebees. Evolution 66, 3765–3777.

