## Supplementary Material for "Bumblebee queens differ in brain morphology but not learning performance across life stages"

**SUPPLEMENT**

**Table S1:** Sample sizes across colonies for the workers and gynes (7-10 day post-eclosion) and natal queens (shaded in grey) and the bees of unknown age that were not included in analyses.

|  | **Learning Assays** | | | | | **Brain imaging** | | |
| --- | --- | --- | --- | --- | --- | --- | --- | --- |
| **Colony** | **workers** | **gynes** | **natal queen** | **unknown-age workers** | **unknown-age gynes** | **workers** | **gynes** | **natal queen** |
| I | 0 | 1 | 0 | 4 | 6 | 0 | 1 | 0 |
| J | 4 | 0 | 0 | 0 | 0 | 1 | 0 | 0 |
| K | 5 | 0 | 1 | 0 | 0 | 2 | 0 | 1 |
| L | 2 | 0 | 0 | 0 | 0 | 0 | 0 | 0 |
| M | 0 | 1 | 0 | 4 | 0 | 0 | 1 | 0 |
| N | 3 | 6 | 0 | 4 | 0 | 2 | 2 | 1 |
| O | 2 | 6 | 0 | 0 | 1 | 1 | 1 | 0 |
| P | 3 | 0 | 0 | 1 | 0 | 0 | 0 | 0 |
| Q | 0 | 0 | 0 | 1 | 0 | 0 | 0 | 0 |
| R | 1 | 0 | 1 | 3 | 0 | 0 | 0 | 1 |
| S | 0 | 0 | 1 | 2 | 0 | 0 | 0 | 1 |
| T | 0 | 0 | 0 | 0 | 0 | 0 | 0 | 0 |
| U | 0 | 0 | 0 | 3 | 0 | 0 | 0 | 0 |
| V | 5 | 4 | 0 | 4 | 0 | 1 | 2 | 0 |
| W | 5 | 8 | 0 | 3 | 20 | 1 | 3 | 0 |
| X | 3 | 3 | 1 | 2 | 4 | 1 | 0 | 1 |
| Y | 1 | 0 | 1 | 3 | 0 | 1 | 0 | 1 |
| B | 0 | 0 | 1 | 2 | 0 | 0 | 0 | 0 |
| C | 0 | 0 | 1 | 1 | 0 | 0 | 0 | 0 |
| **TOTALS** | **34** | **29** | **7** | 34 | 31 | **10** (one damaged) | **10** | **6** |
| **Non-participants** | 1 | 3 | 2 | 3 | 6 | **Total =** | **26** | |
|  | **Total =** | **70** | | Total of unknown age = | 65 |  |  |  |

| **Brain Region** | **Pairwise** | **Slope Comparison** | **Intcept Comparison** |
| --- | --- | --- | --- |
| Total Reconstructed Volume | Queen vs Gyne | t = 2.013, p = 0.155 | Wald X^2^ = 3.642, p = 0.056 |
|  | Queen vs Worker | t = 0.015, p = 0.901 | **Wald X^2^ = 31.299, p < 0.001** |
|  | Gyne vs Worker | t = 2.410, p = 0.120 | **Wald X^2^ = 5.044, p = 0.024** |
| Medulla | Queen vs Gyne | **t = 4.315, p = 0.037** | NA |
|  | Queen vs Worker | t = 0.053, p = 0.817 | Wald X^2^ = 0.5679, p = 0.451 |
|  | Gyne vs Worker | **t = 4.145, p = 0.041** | NA |
| Lobula | Queens vs Gyne | t = 0.591, p = 0.441 | **Wald X^2^ = 4.285, p = 0.038** |
|  | Queen vs Worker | t = 2.181, p = 0.139 | Wald X^2^ = 0.026, p = 0.871 |
|  | Gyne vs Worker | **t = 2.181, p = 0.007** | NA |
| Antennal Lobe | Queen vs Gyne | t = 4.070, p = 0.103 | **Wald X^2^ = 11.71, p < 0.001** |
|  | Queen vs Worker | t = 0.492, p = 0.482 | **Wald X^2^ = 92.76, p <0.001** |
|  | Gyne vs Worker | **t = 6.855, p = 0.008** | NA |
| MB Collar | Queen vs Gyne | t = 0.006, p = 0.936 | Wald X^2^ = 0.001, p = 0.973 |
|  | Queen vs Worker | t = 0.563, p = 0.453 | **Wald X^2^ = 18.509, p < 0.001** |
|  | Gyne vs Worker | t = 0.537, p = 0.463 | **Wald X^2^ = 6.629, p = 0.010** |
| MB Lip | Queen vs Gyne | t = 0.148, p = 0.699 | Wald X^2^ = 0.710, p = 0.399 |
|  | Queen vs Worker | t = 0.833, p = 0.361 | **Wald X^2^ = 19.163, p < 0.001** |
|  | Gyne vs Worker | t = 0.382, p = 0.536 | **Wald X^2^ = 6.168, p = 0.013** |

**Table S2:** Standard major axis (SMA) regression results with ITD as the measure of body size; results do not change from using PC1 as the measure of body size.

**Table S3:** emmeans pairwise comparisons from linear models addressing differences in absolute brain volumes (i.e. where body size is not accounted for). General model structure is: (lm(region ~ type)). Significant differences are bolded.

| **Brain Region** | **Pairwise** | **p-value** | **Test Statistic** |
| --- | --- | --- | --- |
| Total Reconstructed Volume | Queen vs Gyne | 0.177 | t(22) = -1.848 |
|  | Queen vs Worker | **0.002** | t(22) = 3.781 |
|  | Gyne vs Worker | 0.083 | t(22) = 2.260 |
| Medulla | Queen vs Gyne | 0.076 | t(23) = -2.298 |
|  | Queen vs Worker | **<0.001** | t(23) = 5.391 |
|  | Gyne vs Worker | **0.004** | t(23) = 3.572 |
| Lobula | Queens vs Gyne | 0.162 | t(23) = -1.895 |
|  | Queen vs Worker | **<0.001** | t(23) = 4.823 |
|  | Gyne vs Worker | **0.007** | t(23) = 3.382 |
| Antennal Lobe | Queen vs Gyne | **<0.001** | t(22) = -4.488 |
|  | Queen vs Worker | **<0.001** | t(22) = 7.075 |
|  | Gyne vs Worker | **0.014** | t(22) = 3.072 |
| MB Collar | Queen vs Gyne | 0.961 | t(22) = -0.266 |
|  | Queen vs Worker | 0.078 | t(22) = 2.287 |
|  | Gyne vs Worker | 0.073 | t(22) = 2.325 |
| MB Lip | Queen vs Gyne | 0.084 | t(22) = -2.254 |
|  | Queen vs Worker | **0.001** | t(22) = 4.055 |
|  | Gyne vs Worker | 0.109 | t(22) = 2.118 |

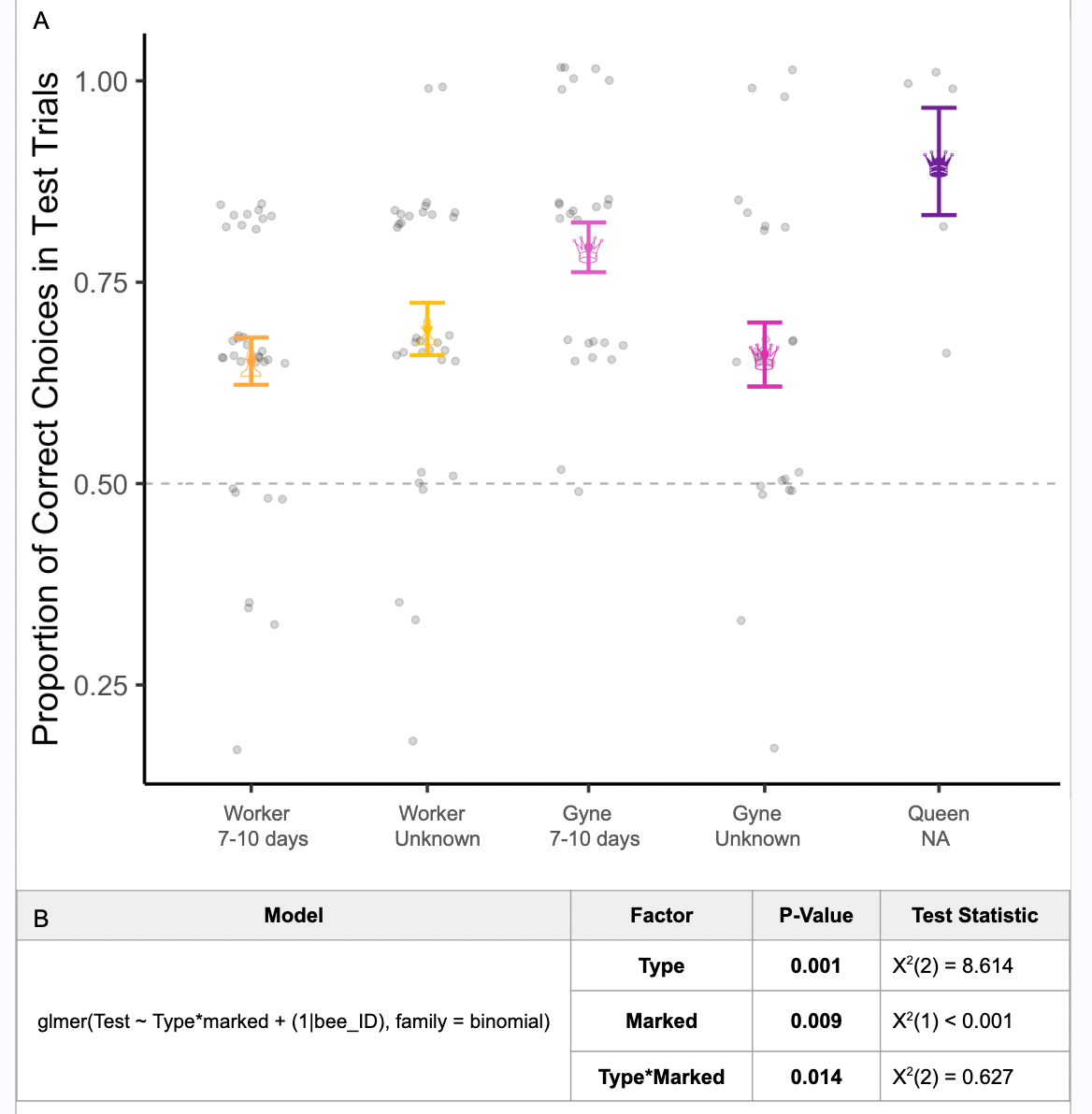

**Figure S1:** In a pilot study we tested an additional set of workers (n = 34) and gynes (n = 31) that were not marked upon eclosion (i.e. bees were of unknown ages) from 15 colonies. These bees were otherwise the same as marked bees: they had no free-flying foraging experience and tested them in the color learning assay, but we found behavioral differences between these bees and our 7-10 day old marked bees. **A:** Test trial performance for the three bee types (queen, gyne, or worker) visualized by if they were marked (7-10 days post eclosion) or were unmarked (unknown age), points are jittered for clarity. **B:** Model output showing that while bee type (queen, gyne or worker) differed in their test trial performance, if they were marked (known vs unknown age) also affected their test trial performance, and this interacted with bee type. Since unknown age bees did not perform similarly to our known age marked bees, we only consider age-matched gynes and workers in our study.

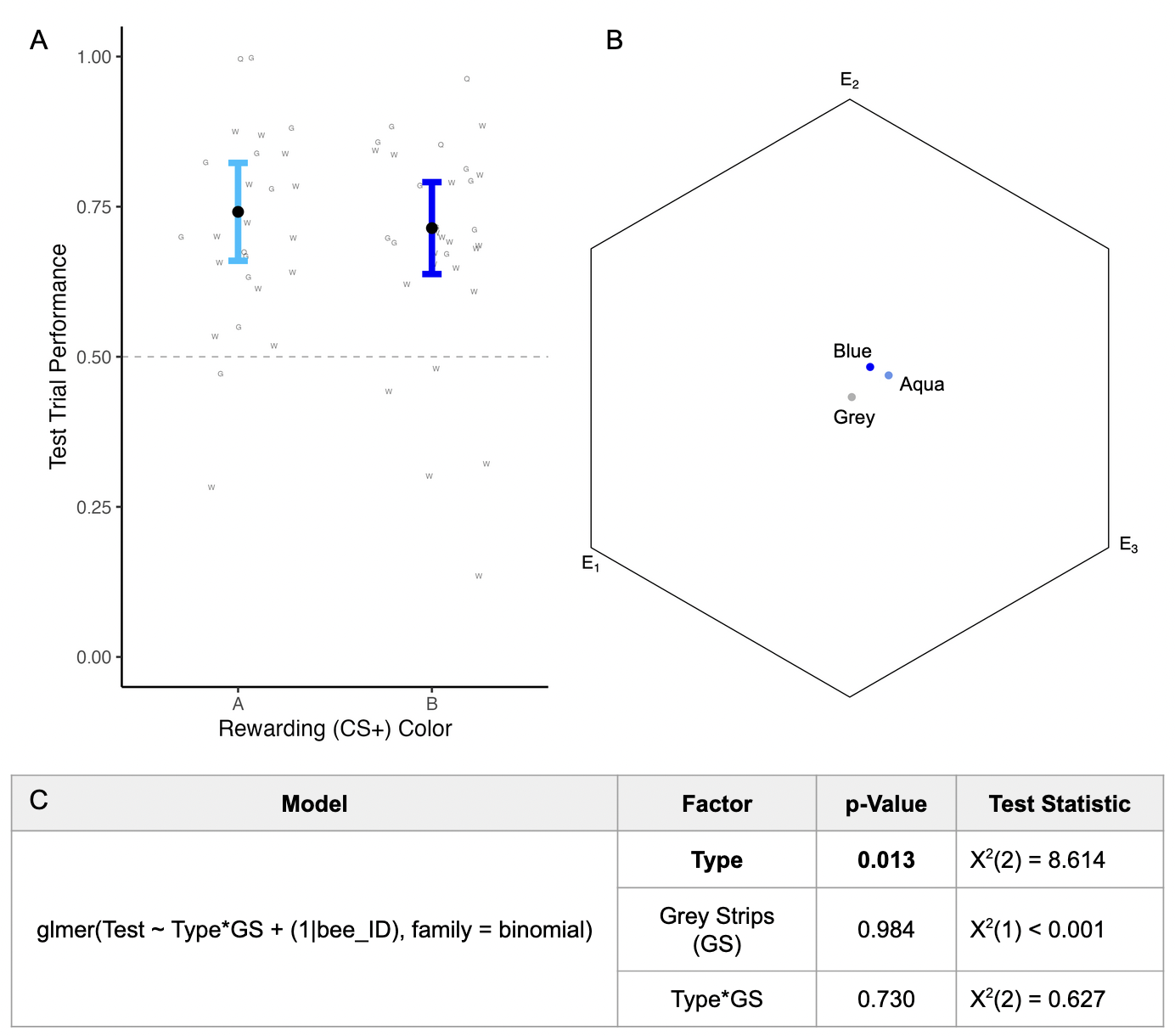

**Figure S2:** **A:** Test trial performance for bees by color trained to as the CS+ (A = aqua, and B = blue); data points refer to the bee type (Q = queen, G = gyne, and W = worker). 29 bees were trained to aqua (Q = 3, G = 12, W = 14) and 35 bees were trained to blue (Q = 2, G = 13, W = 20) **B:** The two blue colors and the grey strips plotted into *Bombus impatiens* color hexagon space, showing that the two blue colors are more chromatically similar to each other than to the grey. **C:** Model output showing that while bee type (queen, gyne, or worker) differed in their test trial performance, presentation of re-motivating grey strips (‘GS’ in model) (0, 1, or 2) did not, and there was no interaction between bee type and grey strip presentation on test trial performance.

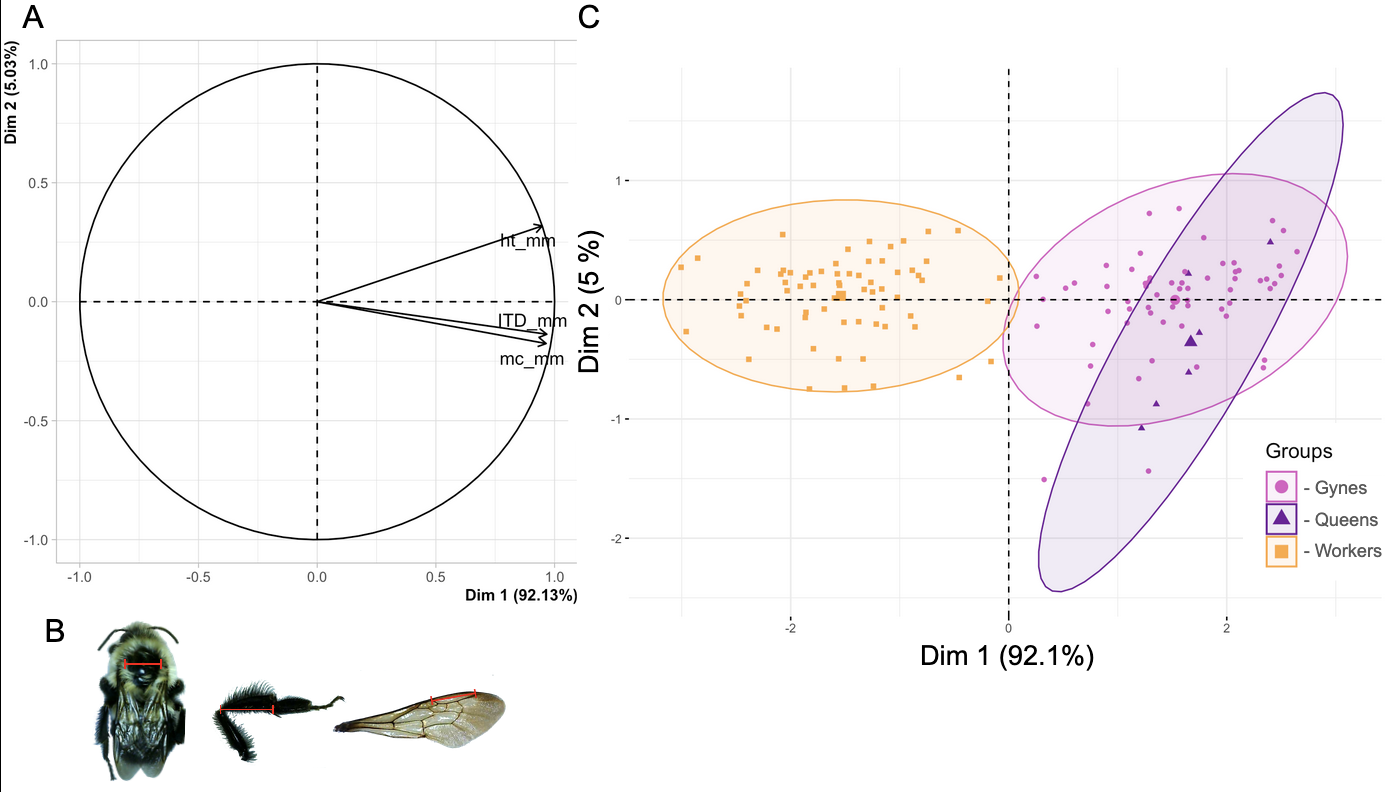

**Figure S3: A:** PCA graph of body size variables, showing that hind tibia length (ht), intertegular distance (ITD), and marginal cell length (mc) are all highly correlated and contribute similarly to PC1. **B:** Diagram showing the three body size metrics (ITD, ht, and mc) **C:** Body size PCA for the three bee types (workers (W), gynes (G), and queens (Q)), showing that PC1 describes 92.1% of the body size variation across our bees.

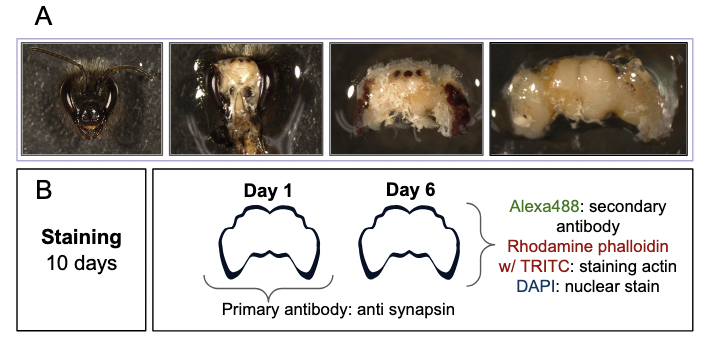

**Fig S4:** **A:** Dissection images showing how the cuticle was removed from the head around the brain in preparation for staining. **B:** Timing of the staining protocol showing the three fluorophores.

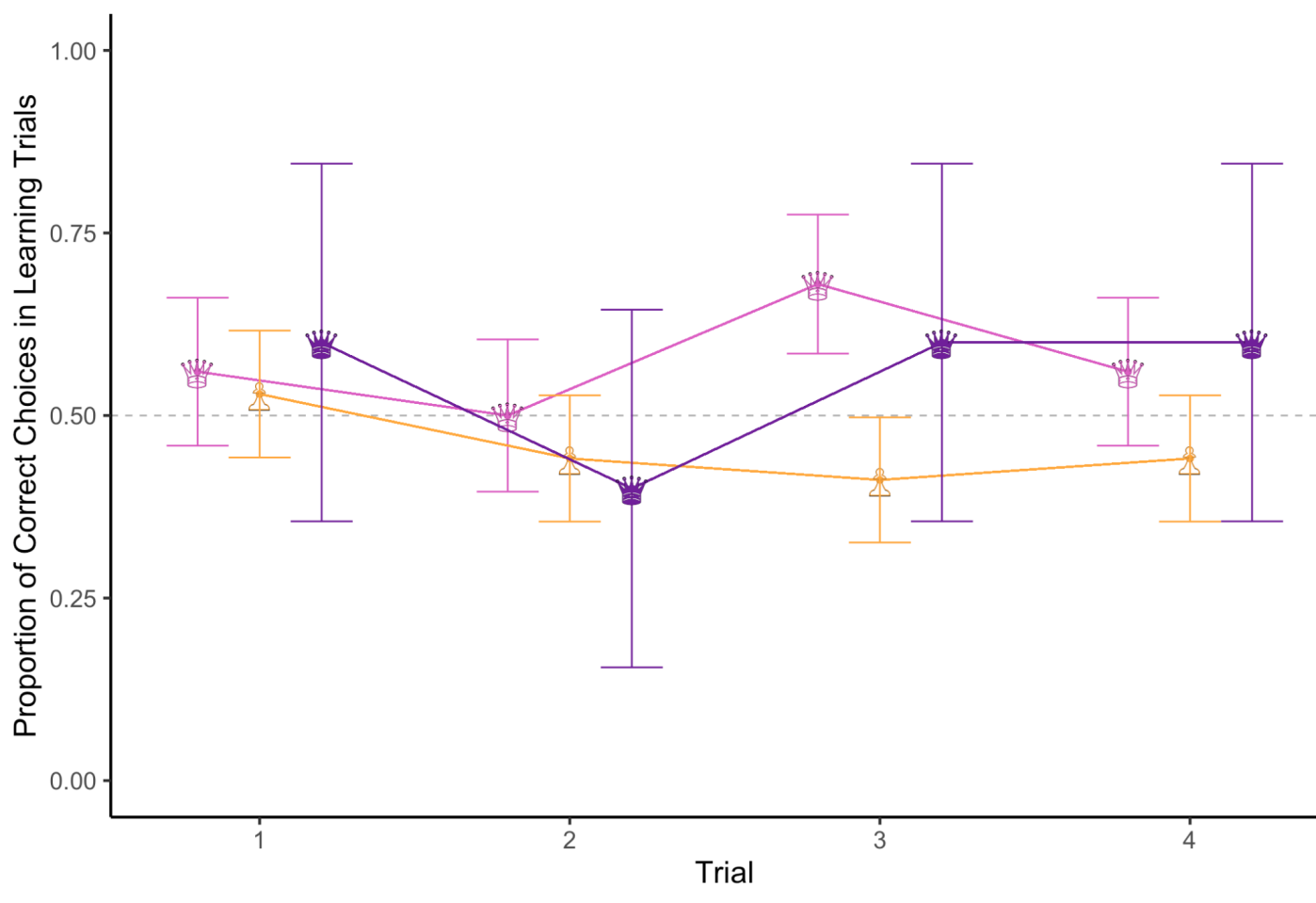

**Figure S5:** Performance of each group (mean +/- SE) over the 4 rewarded learning trials. The gray dotted line represents performance at chance (50%).
